# *In silico* transcriptomic analysis reveals shared molecular signatures and immune-associated pathways between Hashimoto’s thyroiditis and type 2 diabetes with exploratory drug repurposing

**DOI:** 10.64898/2026.02.16.706089

**Authors:** Ovi Sharma, Md. Feroj Ahmed, Dipu Sharma, Apurbo Sharma, Tasfia Noor, Fahim Faysal, Md. Foysal Ahmed, Md. Sanoar Hossain, Md. Al Noman, Md. Abdul Latif, Mohammad Ali, Dewan Mumdood Ahmed, Md. Nurul Haque Mollah

**Affiliations:** Bioinformatics Lab (Dry), Department of Statistics, University of Rajshahi, Rajshahi-6205, Bangladesh; Bachelor of Medicine and Bachelor of Surgery (MBBS), TMSS Medical College, Bogura, Bangladesh; Department of Chemistry, University of Rajshahi, Rajshahi-6205, Bangladesh; Institute of Bangladesh Studies (IBS), University of Rajshahi, Rajshahi-6205, Bangladesh; Department of ICE, Pabna University of Science and Technology, Pabna, Bangladesh; Department of Computer Science & Engineering, Rajshahi University of Engineering & Technology, Rajshahi-6204, Bangladesh

**Keywords:** Hashimoto’s thyroiditis, Type 2 diabetes, shared key genes, shared molecular mechanism, Immune regulation, Drug repurposing

## Abstract

The management of Hashimoto’s thyroiditis (HT), one of the most prevalent autoimmune disorders worldwide, becomes more complex when it coexists with type 2 diabetes (T2D) compared with the management of either disease alone. This complexity may arise from overlapping genetic, metabolic, and immune dysregulation, as well as potential therapeutic conflicts. Although HT and T2D are known to co-occur and share immune and metabolic features, the molecular characteristics underlying these overlaps have not been systematically explored. This study aimed to identify shared gene expression signatures and associated biological pathways between HT and T2D using an in silico, hypothesis-generating approach, and to explore candidate compounds that may be relevant to both conditions. Independent transcriptomic datasets (GSE138198 for control/HT and GSE29231 for control/T2D) were analyzed, leading to the identification of 59 genes that were differentially expressed in both HT and T2D compared with control samples. Protein-protein interaction (PPI) network analysis prioritized five shared key genes (sKGs): CDC42, CD74, FOS, RAC2, and YWHAB. Functional enrichment analysis of these sKGs revealed overlapping biological processes, molecular functions, cellular components, and immune-related signaling pathways, as well as shared regulatory networks involving transcription factors (FOXC1 and HNF4A) and microRNAs (hsa-miR-221-3p and hsa-miR-29a-3p). Immune infiltration analysis demonstrated broadly similar patterns of immune dysregulation in both diseases, providing additional biological context for the observed shared molecular signatures. Finally, an exploratory in silico drug repurposing pipeline incorporating molecular docking, ADMET profiling, drug-likeness assessment, and molecular dynamics simulations prioritized three candidate compounds: gliquidone, oleanolic acid, and glipizide for further investigation. Overall, this study provides a hypothesis-generating framework highlighting shared molecular features between HT and T2D, which may inform future experimental validation and clinical research.

**Author Summary:** In this work, we wanted to better understand why Hashimoto’s thyroiditis, an autoimmune condition that affects the thyroid gland, is often seen in people who also have type 2 diabetes. Treating patients who live with both conditions can be difficult, and we were interested in finding out whether they share common biological causes. To do this, we examined genetic data from individuals with each disease and looked for patterns that appeared in both groups. We discovered several genes that seem to act in similar ways in the two conditions, particularly genes linked to immune system activity and associated pathways. This finding suggests that shared molecular signatures and immune-associated pathways may play a role in the development of both diseases. We also explored how these shared genetic features influence larger biological processes and immune responses. The similarities we found support the idea that the two diseases may be connected through related biological pathways. In addition, we used computer-based screening methods to identify existing drugs that might influence these shared pathways. While these results need further testing, we hope our findings help open new directions for research and eventually contribute to better care for patients affected by both conditions.

## Introduction

Hashimoto’s Thyroiditis (HT) is a common autoimmune condition characterized by chronic inflammation of the thyroid gland and the production of autoantibodies, often leading to hypothyroidism[1]. Clinically, it presents with thyroid enlargement, fatigue, cold intolerance, and in some cases, an early transient hyperthyroid phase[2]. HT affects approximately 5-10% of the global population, with significantly higher prevalence in women and older adults[3]. Cohort studies such as the Whickham survey and Rotterdam study have reported 13–16% thyroid antibody positivity in adult women, with increased risk of developing overt hypothyroidism over time[4,5]. The pathogenesis of HT involves complex interactions among genetic susceptibility, environmental triggers, and immune dysregulation [6,7]. On a different note, Type 2 diabetes (T2D) is a long-lasting metabolic illness marked by the body’s reduced sensitivity to insulin and difficulties in controlling blood sugar levels. As reported by the Global Burden of Disease Study, T2D cases rose from 211 million in 1990 to over 463 million in 2019, with deaths increasing from 0.9 million to 1.5 million during the same period[8,9]. The age-standardized prevalence rate increased by more than 40% globally [10]. DALYs due to T2D doubled from 25 million to over 50 million, indicating both mortality and severe disability[11,12]. Clinical observations suggest that thyroid dysfunction is relatively common among individuals with T2D[13,14]. Numerous studies have revealed that thyroid dysfunction, particularly subclinical hypothyroidism, is independently associated with insulin resistance and impaired glucose metabolism, both critical features in T2D pathogenesis[15,16]. A large cross-sectional study found that up to 23% of T2D patients had underlying thyroid dysfunction, primarily hypothyroidism. Further, anti-thyroid peroxidase (TPO) antibody positivity has been linked with poor glycemic control and increased peripheral artery disease in diabetic patients[17]. Thyroid hormones significantly influence glucose homeostasis, hepatic gluconeogenesis, and pancreatic β-cell function[18]. Reduced thyroid activity can impair insulin sensitivity and lipid metabolism, thereby contributing to hyperglycemia and weight gain, which are recognized contributors to T2D progression[19]. **Figure 1** illustrates how HT and T2D are interconnected. The schematic draws on data from earlier studies to outline the principal pathways by which thyroid autoimmunity may influence metabolic dysfunction in T2D.

**Figure 1.**
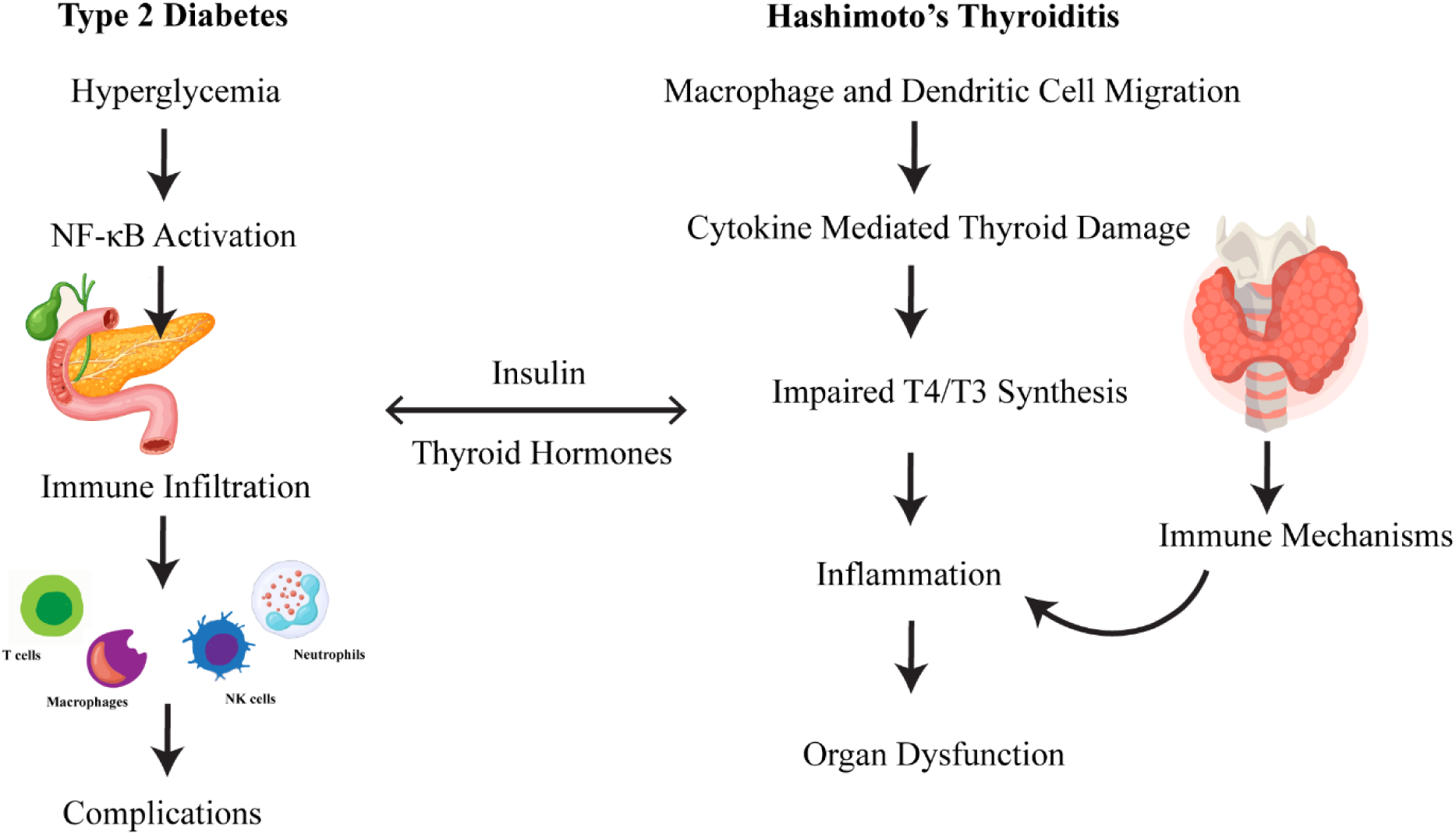
A schematic diagram showing the link between HT and T2D.

Despite increasing epidemiological and clinical evidence of their coexistence, the molecular commonalities between HT and T2D remain incompletely characterized [20,21]. Previous studies have independently identified disease-specific key genes and pathways for T2D [22,23] and HT[24,25]. However, systematic identification of shared gene expression signatures across these two conditions using transcriptomic data has been limited. Identifying such shared molecular features may provide insight into common biological processes without implying direct causal interaction between the diseases. In addition, drug repurposing strategies have gained attention as cost-effective approaches for identifying candidate compounds that may target shared molecular pathways across diseases. While established therapies exist for T2D, including metformin [26] and pioglitazone[27], treatment for HT primarily involves hormone replacement therapy such as levothyroxine[28]. No studies to date have systematically explored potential candidate compounds that may theoretically interact with shared molecular targets in both conditions using computational approaches. Such analyses may help prioritize compounds for future experimental investigation, although clinical applicability requires further validation [29–31]. Therefore, in this study, we performed a comparative transcriptomic analysis of independent HT and T2D datasets to identify shared differentially expressed genes (sDEGs) and shared key genes (sKGs) [32–34].

We further explored associated functional pathways, regulatory networks, immune cell infiltration patterns, and conducted an exploratory in-silico drug prioritization analysis. Our objective was to provide a hypothesis-generating framework describing shared molecular signatures between HT and T2D. Although the findings require validation through mechanistic wet-lab experiments and rigorously designed clinical trials, the computational prioritization approach presented here may facilitate more efficient experimental design by narrowing candidate targets, thereby potentially reducing time, cost, and laboratory burden.

**Figure 2.**
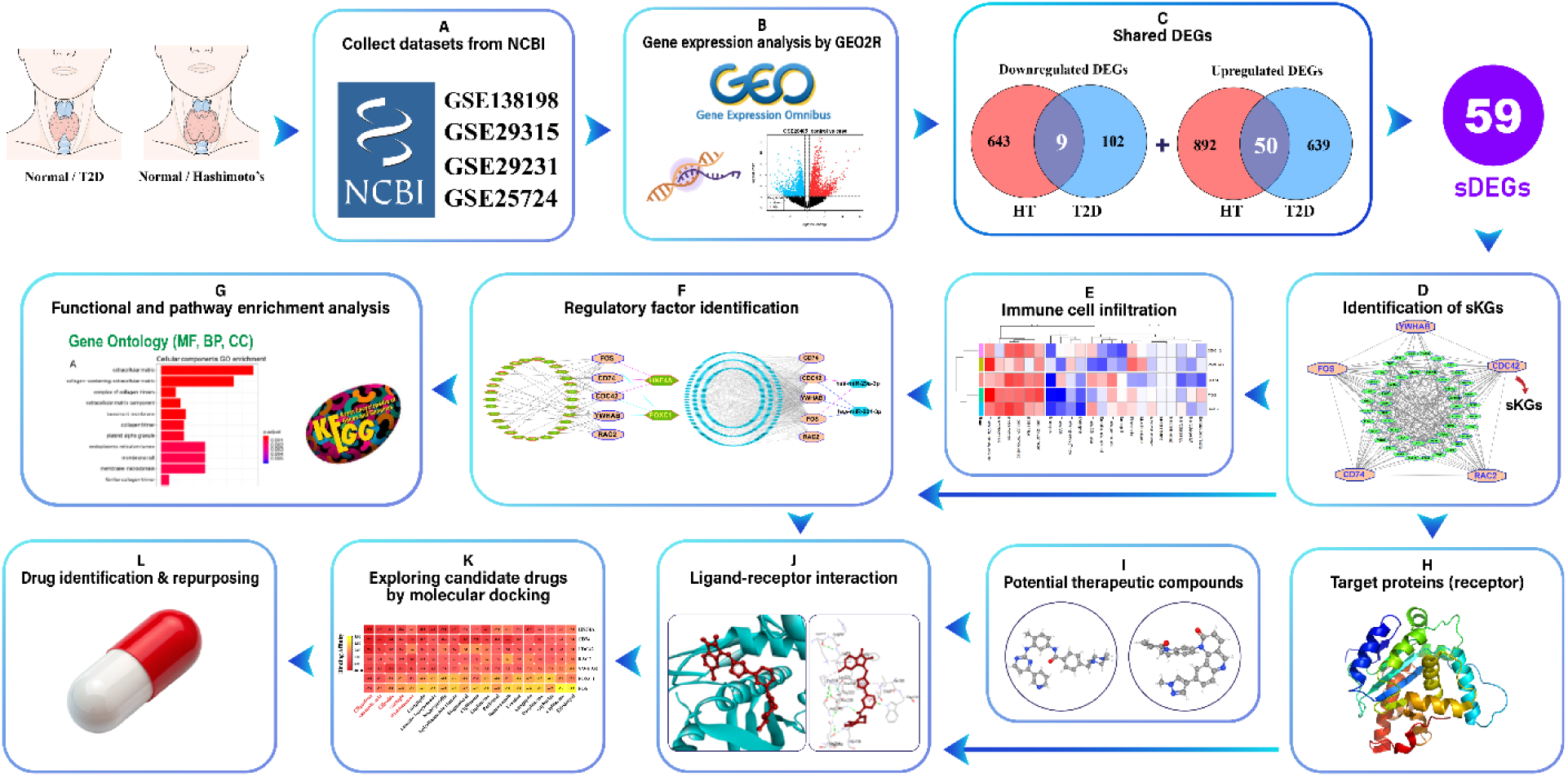
A graphical representation of the entire workflow.

## Results

### Identification of Differentially Expressed Genes (DEGs)

DEGs were identified separately within each dataset using the LIMMA package in R. DEGs were selected based on a cut-off of adjusted P-values < 0.05 and |Log2FC| > 1. For HT, analysis of the NCBI dataset GSE138198 revealed 942 upregulated and 647 downregulated DEGs. In T2D patients, analysis of the NCBI dataset GSE29231 identified 689 upregulated and 106 downregulated DEGs (**Table S2 & Table S4**).

### Identification of shared DEGs (sDEGs)

As reported in the previous section, analysis of the transcriptomic dataset for HT revealed 892 upregulated and 643 downregulated DEGs. Similarly, 102 downregulated and 639 upregulated DEGs for T2D were identified from the transcriptomics dataset. Next, we identified 50 upregulated and 9 downregulated sDEGs between HT and T2D (**Tables S3 & Table S4),** yielding a total of 59 sDEGs across both conditions. These sDEGs represent genes exhibiting concordant differential expression patterns in both conditions compared with controls.

**Figure 3.**
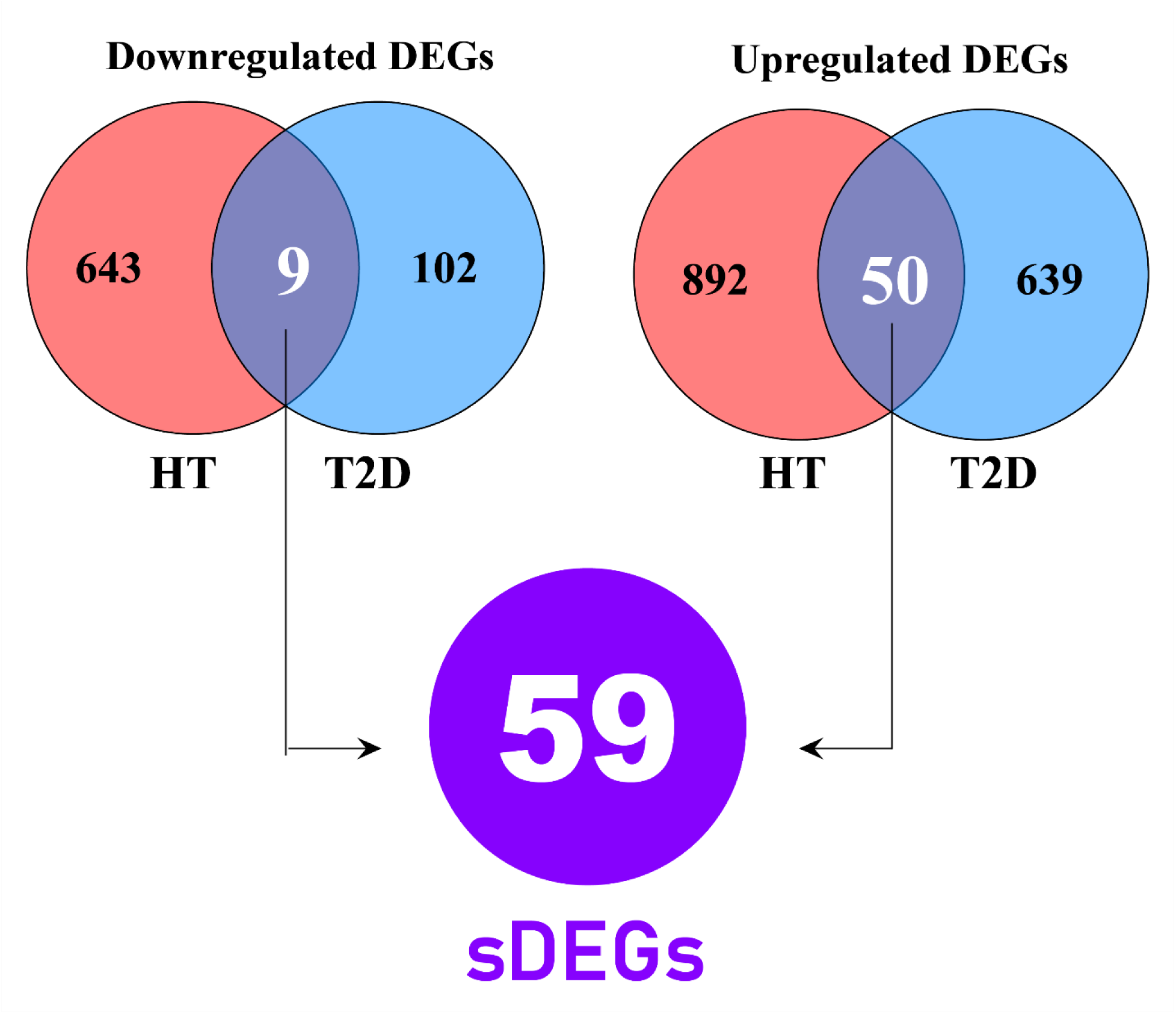
Identification of sDEGs between HT and T2D. The left Venn diagram shows down-regulated DEGs (HT: 643, T2D: 102, overlap: 9), and the right Venn diagram shows up-regulated DEGs (HT: 892, T2D: 639, overlap: 50), totaling 59 sDEGs.

### Identification of shared key genes (sKGs) from sDEGs

STRING was employed to generate the protein-protein interaction (PPI) network of sDEGs. Five topological analysis methods-Betweenness, Closeness, Degree, MCC, and EPC were employed on the PPI network to identify genes crucial for network stability and biological function. We chose the top 5 sKGs: CDC42, CD74, RAC2, FOS, and YWHAB **(Table S5)**. These genes were prioritized using network topology-based analyses and should not be interpreted as definitive causal drivers of disease without further experimental validation.

**Figure 4.**
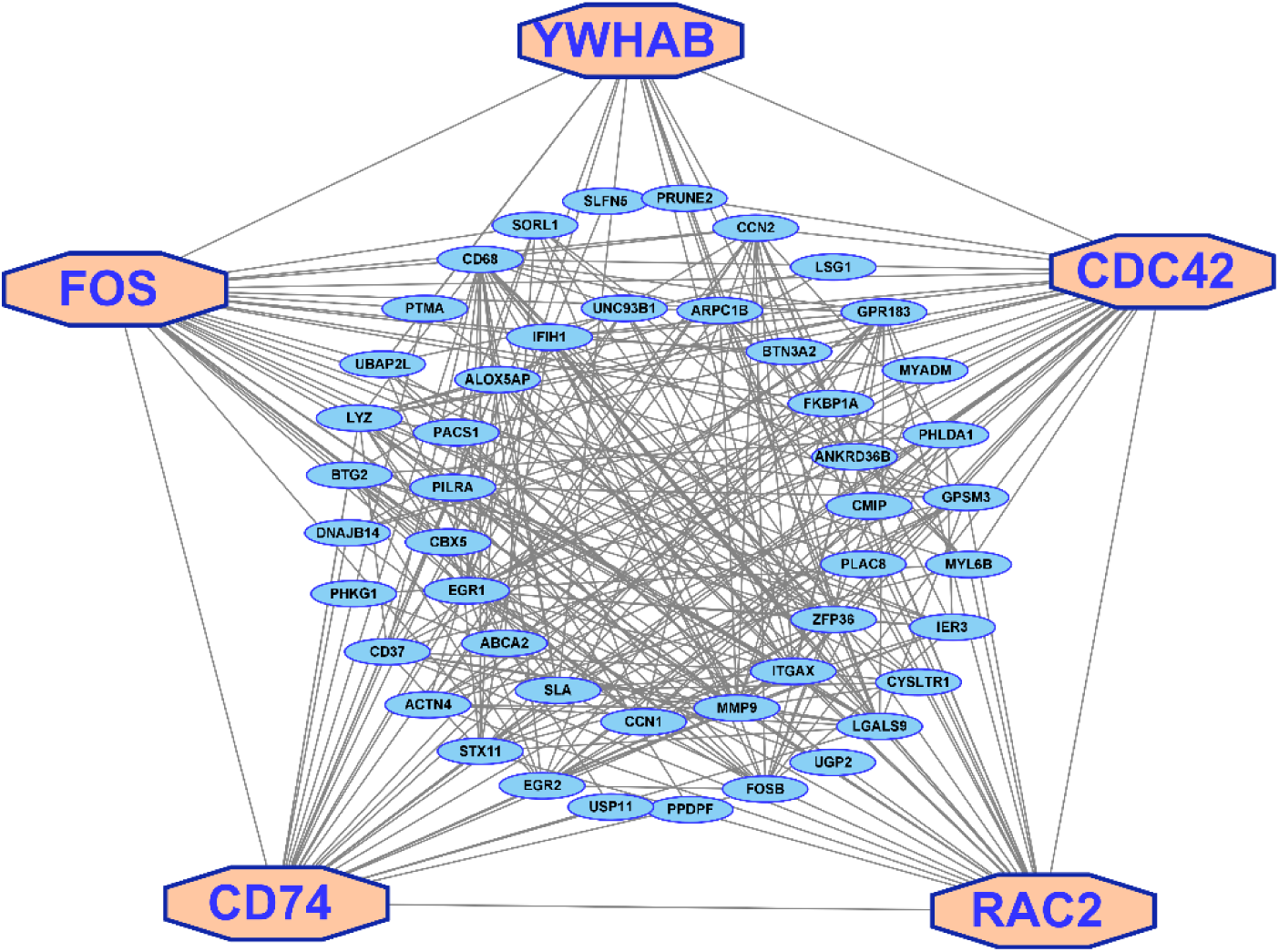
PPI network of sDEGs for identification of shared key-genes (sKGs).

### *In-silico* validation of sKGs using independent transcriptomic datasets

Differential expression of the sKGs across disease and control groups was analyzed through box plot representation. Independent gene expression datasets were retrieved from the NCBI GEO database: GSE29315, which includes 6 Hashimoto’s thyroiditis and 8 control samples **(Figure 5)**, and GSE25724, comprising 6 T2D and 7 control samples **(Figure 6)**. Using the ‘ggpubr’ R package, we confirmed the expression of the key genes. As shown in Figure 5, 6 all five sKGs-CDC42, FOS, CD74, RAC2, and YWHAB were found to be upregulated in both disease conditions, supporting our proposed findings.

**Figure 5.**
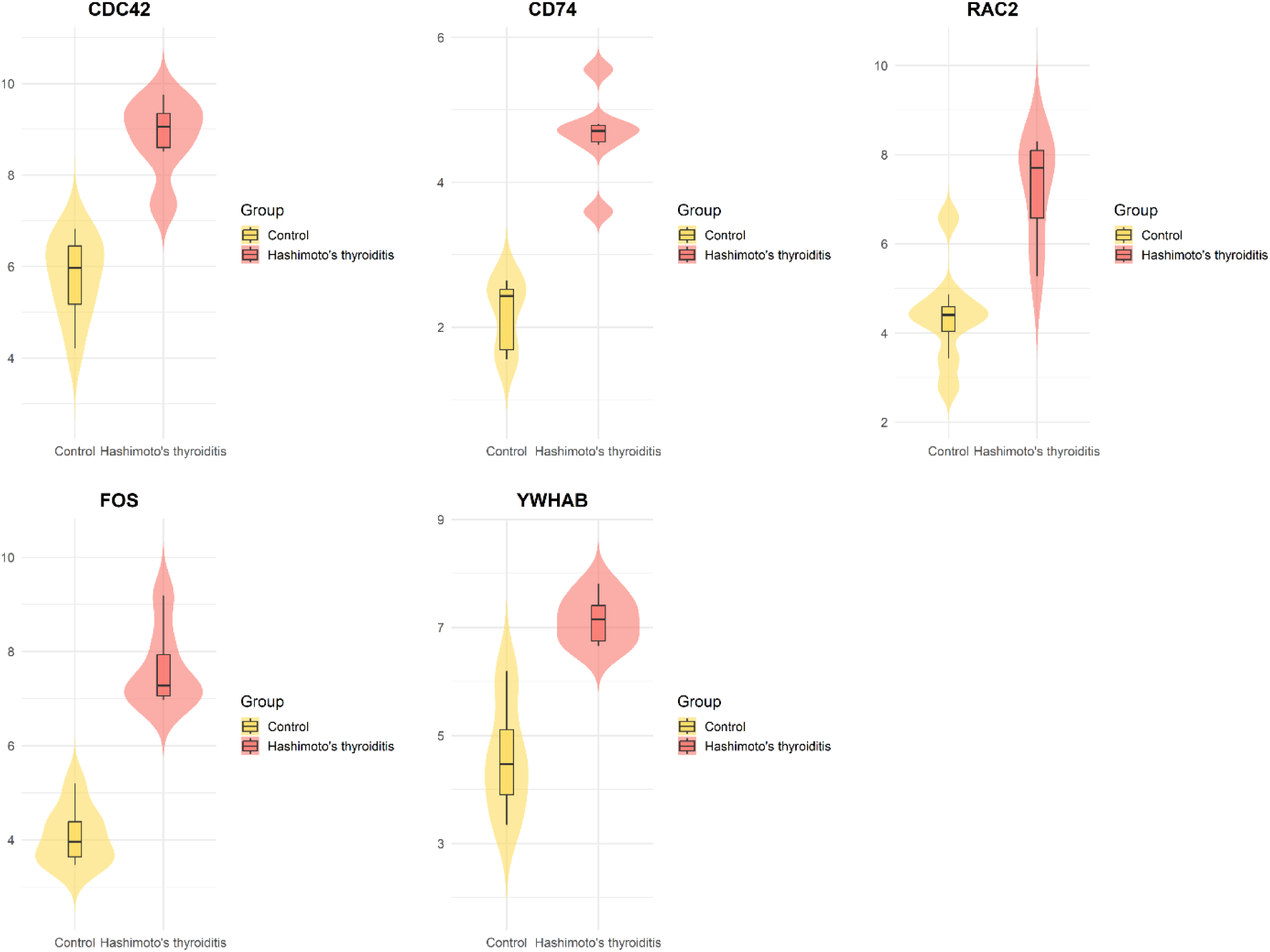
Violin plots displaying the gene expression patterns of sKGs (CDC42, CD74, RAC2, FOS, and YWHAB) between control and HT groups in the dataset with NCBI accession ID GSE-29315. Each plot compares the expression patterns between HT and control samples.

**Figure 6.**
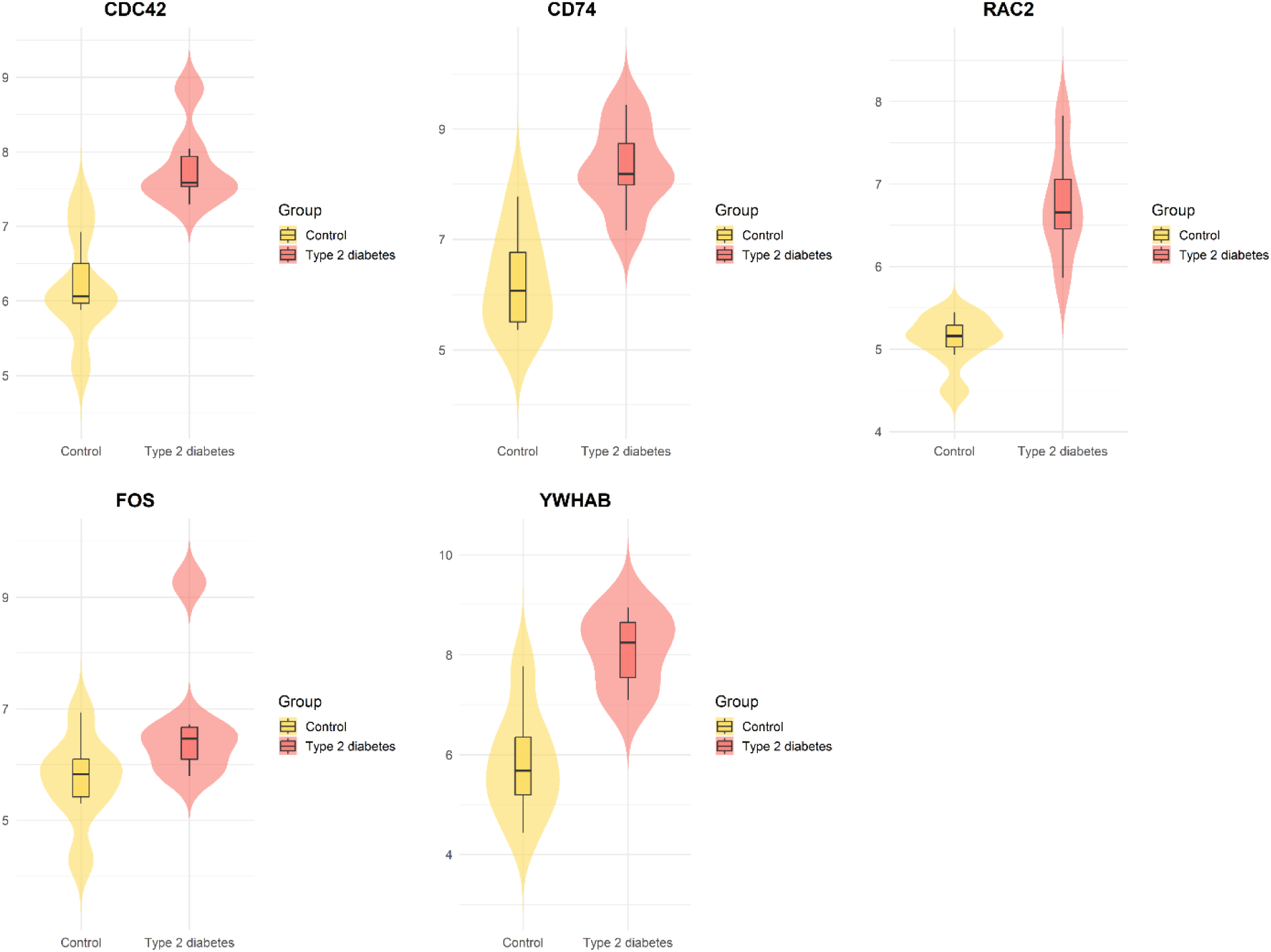
Violin plots displaying the gene expression patterns of sKGs (CDC42, CD74, RAC2, FOS, and YWHAB) between control and T2D groups in the dataset with NCBI accession ID GSE-25724. Each plot compares the expression pattern between T2D and control samples.

### Elucidating shared pathogenetic mechanisms

Shared key genes were used to investigate regulatory factors, biological processes, molecular functions, cellular components, and pathways to reveal common pathogenetic mechanisms between HT and T2D, as detailed below.

### The regulatory network analysis of sKGs

To identify both transcriptional and post-transcriptional regulators of the sKGs, we investigated the regulatory networks involving transcription factors (TFs) and microRNAs (miRNAs) associated with sKGs **(Figure 7)**. We initially selected FOXC1 and HNF4A as the top transcriptional regulators of sKGs, applying cutoffs of degree ≥ 2 and betweenness centrality ≥ 50.61. Applying the same topological criteria, degree≥ 2 and betweenness centrality ≥ 1008. Hsa-miR-221-3p and Hsa-miR-29a-3p were found to be the highest-ranked microRNAs linked to the sKGs. These findings represent predicted regulatory associations derived from curated databases and network topology and do not establish direct regulatory or causal relationships.

**Figure 7.**
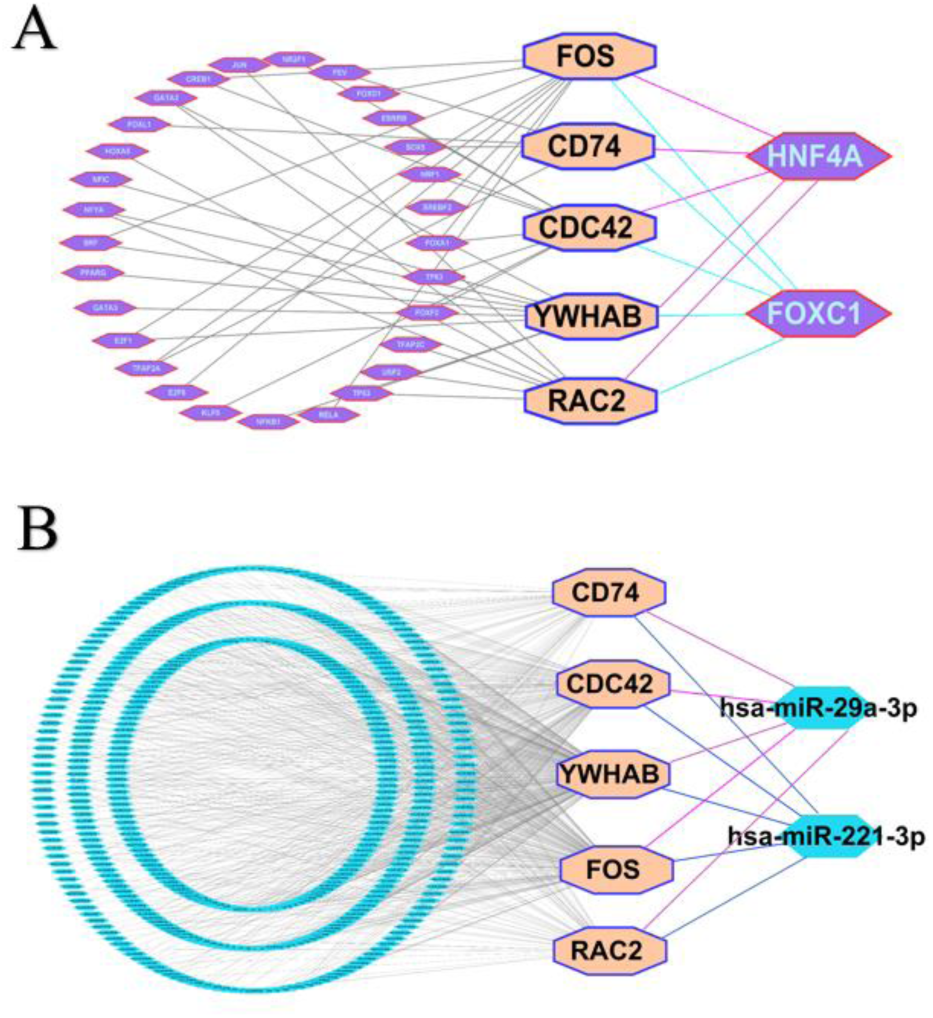
The interaction network visualizes the relationships between (A) TFs and sKGs and (B) miRNAs and sKGs, where larger octagonal nodes indicate sKGs in both A and B. Two larger hexagonal nodes in A and two larger hexagonal nodes in B, indicated by the top-ranked TFs and miRNAs, respectively, represent the key regulators of sKGs.

### Functional enrichment analysis of sKGs

Gene Ontology (GO) and KEGG pathway enrichment analyses were performed for the five sKGs using Enrichr[35]. After multiple testing correction using the Benjamini–Hochberg false discovery rate (FDR), six biological process (BP) terms, five cellular component (CC) terms, five molecular function (MF) terms, and four KEGG pathways were identified as significantly enriched. As summarized in Table 2 and Tables S6–S8, these enriched terms represent shared functional themes and pathway associations observed in both HT and T2D. These results provide descriptive functional context for the sKGs rather than establishing disease-specific mechanisms or validated therapeutic targets.

**Table 1:** Detailed information about the transcriptomic datasets analyzed in this study.

| Accession IDs for datasets | Disease name | Country | Platform | Case | Control |
| --- | --- | --- | --- | --- | --- |
| GSE138198 | HT | Saudi Arabia | GPL6244 [HuGene-1_0-st] Affymetrix Human Gene 1.0 ST Array [transcript (gene) version]. | 13 | 3 |
| GSE29315 | HT | Belgium | GPL8300 [HG_U95Av2] Affymetrix Human Genome U95 Version 2 Array. | 6 | 8 |
| GSE29231 | T2D | India | GPL6947illumina HumanHT-12 V3.0 expression beadchip. | 12 | 12 |
| GSE25724 | T2D | Italy | GPL96 [HG-U133A] Affymetrix Human Genome U133A Array. | 6 | 7 |

**Table 2.** Significant enrichment (P < .05) of GO terms and KEGG pathways associated with sKGs was observed in Enrich.

| **Biological Process (BP)** |  |  |  |
| --- | --- | --- | --- |
| **Annotation ID** | **Terms** | **Adjusted *P*-Value** | **Associated sKGs** |
| GO:0032956 | Regulation of Actin Cytoskeleton Organization | 2.65E-04 | CDC42, RAC2 |
| GO:0090023 | Positive Regulation of Neutrophil Chemotaxis | 6.32E-04 | CD74, RAC2 |
| GO:0032675 | Regulation of Interleukin-6 Production | 0.02818 | CD74 |
| GO:0045944 | Positive Regulation of Transcription by RNA Polymerase II | 0.030655 | YWHAB, FOS |
| GO:0034599 | Cellular Response to Oxidative Stress | 0.038598 | FOS |
| GO:0045893 | Positive Regulation of DNA-templated Transcription | 0.01622 | CD74, YWHAB, FOS |
| **Molecular Function (MF)** |  |  |  |
| GO:0003924 | GTPase Activity | 0.001716 | CDC42, RAC2 |
| GO:0004896 | Cytokine Receptor Activity | 0.036483 | CD74 |
| GO:0003924 | GTPase Activity | 0.009742 | CDC42, RAC2 |
| GO:0001046 | Core Promoter Sequence-Specific DNA Binding | 0.020507 | FOS |
| GO:0019212 | Phosphatase Inhibitor Activity | 0.020089 | YWHAB |
| **Cellular Component (CC)** |  |  |  |
| GO:0005925 | Focal Adhesion | 6.98E-05 | CDC42, YWHAB, RAC2 |
| GO:0005789 | Endoplasmic Reticulum Membrane | 0.008504 | CDC42, CD74, RAC2 |
| GO:0098588 | Bounding Membrane of Organelle | 0.04361 | CDC42, CD74 |
| GO:0000139 | Golgi Membrane | 0.029642 | CDC42, CD74 |
| GO:0098858 | Actin-Based Cell Projection | 0.043618 | CDC42 |
| **KEGG Pathway** |  |  |  |
| hsa04010 | MAPK signaling pathway | 0.001276 | CDC42, RAC2, FOS |
| hsa04612 | Antigen processing and presentation | 0.01935 | CD74 |
| hsa04657 | IL-17 signaling pathway | 0.045767 | FOS |
| hsa04390 | Hippo signaling pathway | 0.050422 | YWHAB |

### Correlation analysis between sKGs and immune cell infiltration

We applied the CIBERSORT algorithm to the GSE138198 and GSE29231 datasets in order to estimate the proportions of 22 immune cell types from bulk transcriptomic data (shown in Figures 8A, 8A). Only samples meeting the CIBERSORT deconvolution significance threshold (P < 0.05) were included in downstream analyses. In HT samples, we saw elevated memory B cells, activated memory CD4⁺ T cells, follicular helper T cells, and activated dendritic cells, and mast cells, while T cells CD4⁺ naive and activated NK cells were reduced (**Figure 8A**). In T2D, memory B cells, activated memory CD4⁺ T cells, follicular helper T cells, M0 macrophages, and activated NK cells were increased; in contrast, levels of CD8⁺ T cells, resting memory CD4⁺ T cells, resting NK cells, M1 macrophages, and neutrophils declined (**Figure 8B**). Importantly, both diseases shared certain immune shifts: memory B cells, activated memory CD4⁺ T cells, and follicular helper T cells were consistently up, while resting memory CD4⁺ T cells and M2 macrophages were down. Correlation analysis demonstrated significant positive associations between sKGs and the elevated immune cell types, and negative associations with the reduced types (P < 0.05) (**Figure 9C, D**). These patterns suggest interactions between hub genes and specific immune populations may contribute to the progression of T2D in HT.

**Figure 8.**
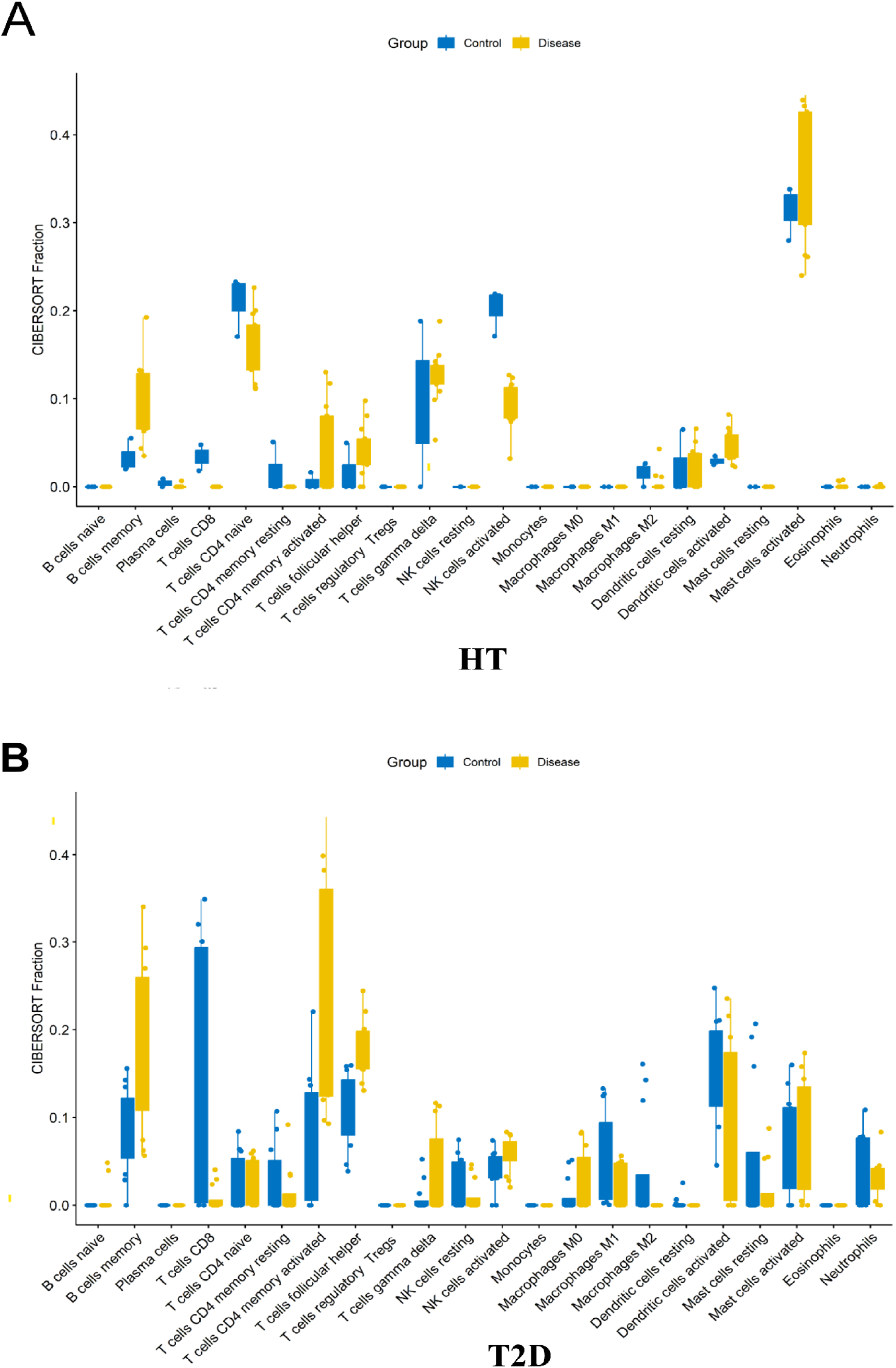
Comparison of immune cell proportions between (A) HT vs. controls, and (B) T2D vs. controls.

**Figure 9.**
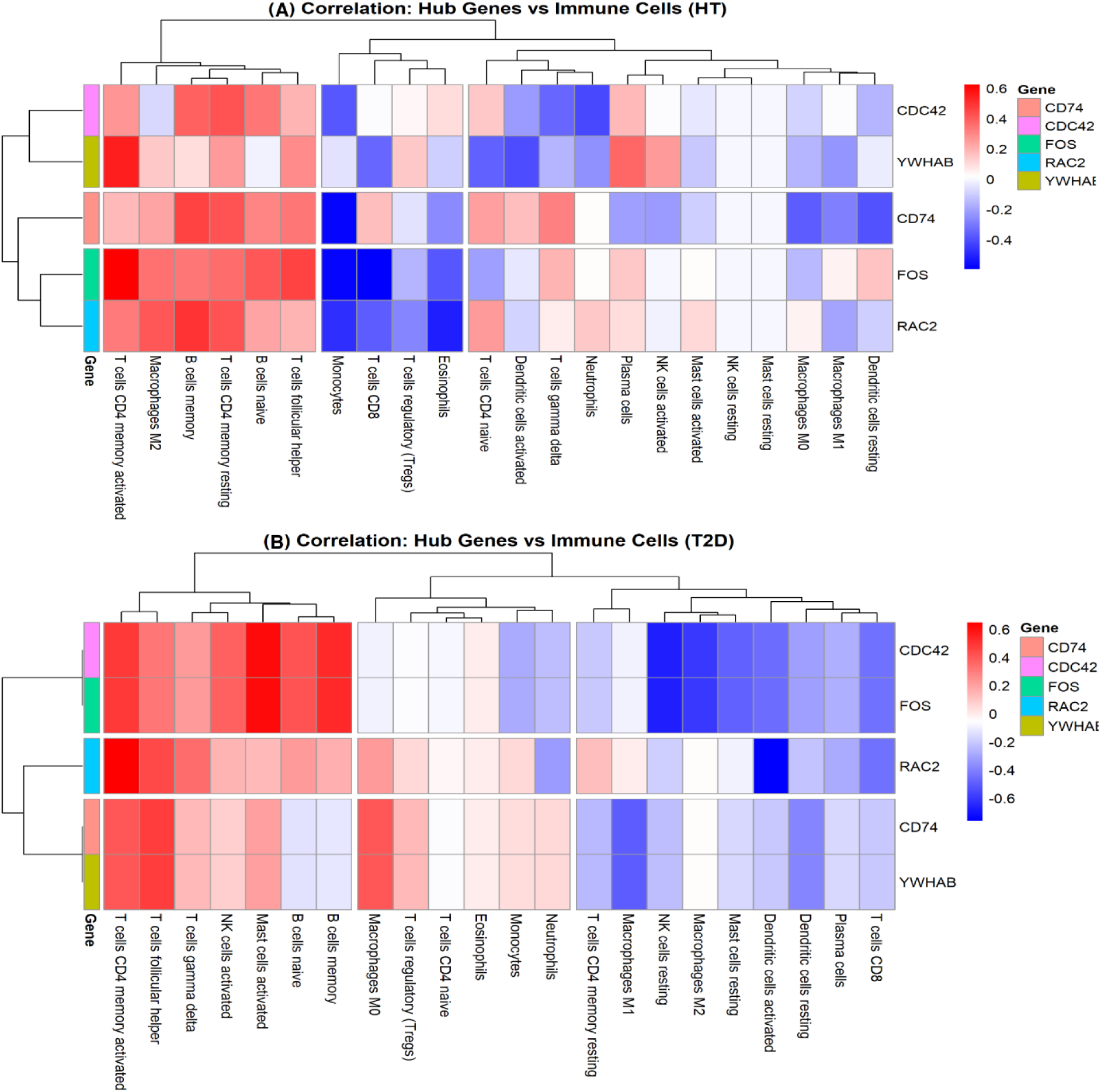
Heatmaps showing correlations between sKGs and immune cells for the datasets with NCBI accession ID (A) GSE138198 for HT and (B) GSE29231 for T2D.

### Druggability Assessment of sKGs

**Table 3.** Binding Pocket Identification and Druggability Assessment of sKGs.

| Gene | Best Pocket site | Volume (Å<sup>3</sup>) | Surface (Å<sup>2</sup>) | Depth(Å) | Hydrophobicity ratio | Druggability Score |
| --- | --- | --- | --- | --- | --- | --- |
| **CDC42** | pocket (P<sub>0</sub>) | 1182.08 | 1393.98 | 26.11 | 0.45 | 0.81 |
| **CD74** | pocket (P<sub>0</sub>) | 1175.81 | 705.23 | 21.47 | 0.42 | 0.82 |
| **RAC2** | pocket (P<sub>0</sub>) | 635.01 | 882.50 | 17.77 | 0.51 | 0.80 |
| **FOS** | pocket (P<sub>0</sub>) | 417.34 | 612.17 | 17.79 | 0.32 | 0.75 |
| **YWHAB** | pocket (P<sub>0</sub>) | 518.22 | 647.77 | 16.72 | 0.54 | 0.82 |
| **FOXC1** | pocket (P<sub>0</sub>) | 4361.56 | 6519.69 | 40.69 | 0.52 | 0.80 |
| **HNF4A** | pocket (P<sub>0</sub>) | 1354.88 | 1191.22 | 30.93 | 0.30 | 0.81 |

Pocket analysis identified several promising druggable binding sites across the selected proteins. CDC42 and CD74 showed large and deep pockets with high druggability scores (CDC42: 1182.08 Å³, depth 26.11 Å, score 0.81; CD74: 1175.81 Å³, depth 21.47 Å, score 0.82), suggesting strong potential for stable ligand binding. RAC2 (635.01 Å³, depth 17.77 Å, score 0.80) and YWHAB (518.22 Å³, depth 16.72 Å, score 0.82) also displayed favorable binding environments with balanced hydrophobic characteristics that can support efficient docking interactions. FOS showed a comparatively smaller pocket (417.34 Å³, depth 17.79 Å, score 0.75), which may be more suitable for selective or fragment-sized ligands. Interestingly, FOXC1 presented a very large and deep cavity (4361.56 Å³, depth 40.69 Å, score 0.80), indicating a flexible binding interface capable of accommodating diverse ligands, while HNF4A also demonstrated a large pocket (1354.88 Å³, depth 30.93 Å, score 0.81) suitable for stable ligand interaction and regulatory targeting. Overall, these pocket characteristics, including volume, depth, hydrophobicity, and druggability, play a crucial role in improving docking accuracy, ligand accessibility, and binding stability, highlighting their importance in structure-based drug discovery. All residues forming the predicted binding pockets are listed in **Table S09**.

### SKGs - guided drug repurposing

To identify compounds that may interact with the sKGs proteins, we performed an exploratory in-silico screening analysis. Molecular docking was used to assess interactions with the sKGs, followed by evaluation of drug-likeness, pharmacokinetic properties, toxicity, and molecular dynamics (MD) simulations to examine complex stability. These analyses were conducted to prioritize promising candidates for further study.

### Molecular docking

We selected five shared key genes (sKGs) and two associated transcription factors as receptor targets for exploratory docking analysis. High-quality three-dimensional protein structures were obtained for four receptors (CDC42, CD74, YWHAB, and HNF4A) from the Protein Data Bank (PDB IDs: 1E0A, 1IIE, 6GN8, and 3FS1), while predicted structures for FOS, RAC2, and FOXC1 were retrieved from AlphaFold (UniProt IDs: AF_P01100_F1, AF_P15153_F1, and AF_Q12948_F1). Candidate compounds were docked against these targets, and binding affinity scores (BAS) were calculated, with more negative values indicating stronger predicted interactions. Compounds were ranked based on aggregated binding affinity values across receptors. Among the tested molecules, gliquidone, oleanolic acid, and glipizide demonstrated average BAS values below −7 kcal/mol, suggesting favorable predicted binding interactions. Naringin and wedelolactone also exhibited comparable docking profiles. These compounds were therefore prioritized for further computational evaluation (**Figure 10A**).

**Figure 10.**
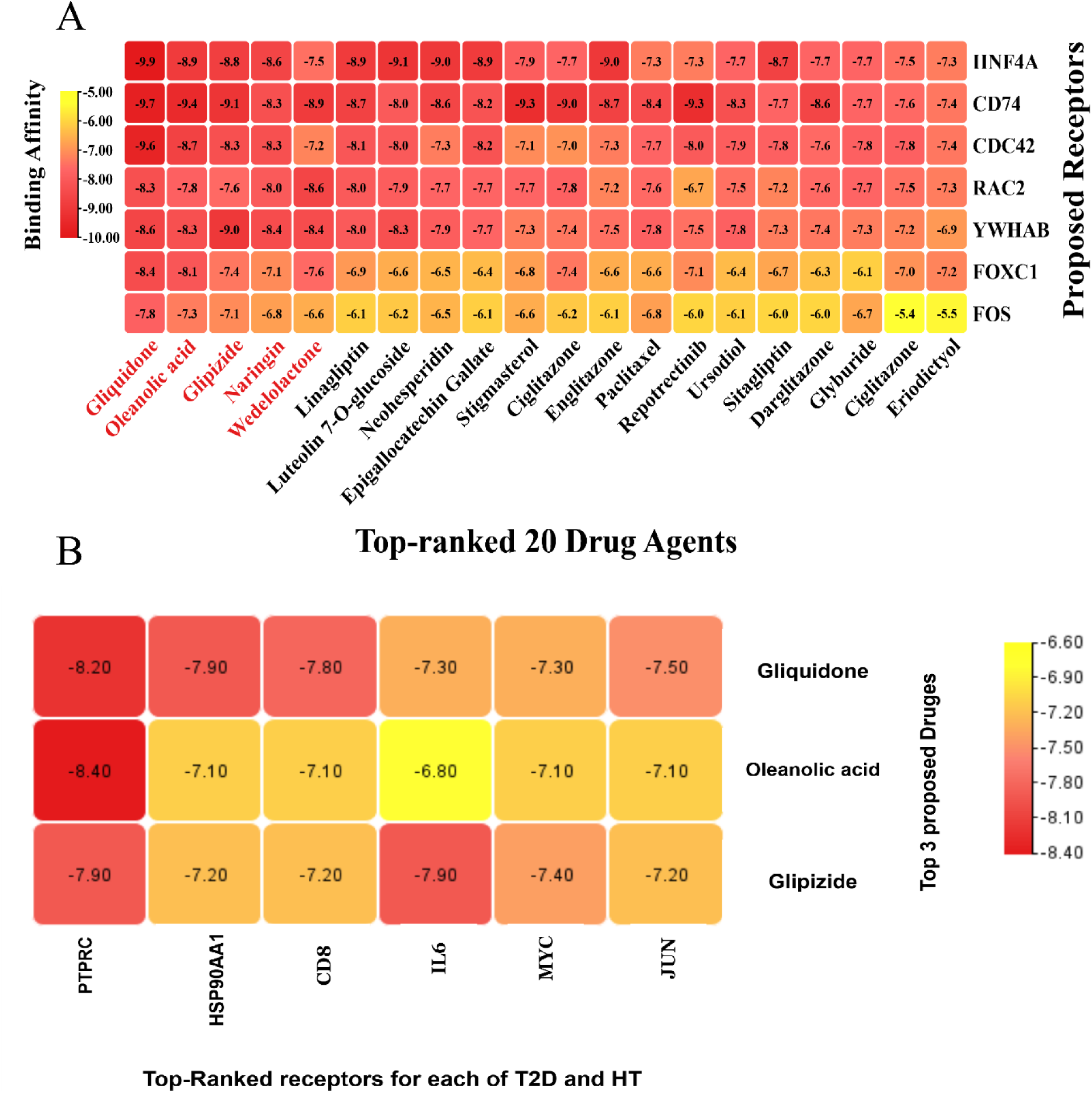
Drug–target binding affinity matrices. (A) The X-axis represents the top 20 ranked drug candidates, while the Y-axis denotes the predicted receptor proteins. (B) On the X-axis, we plot the three highest-ranked both HT- and T2D-causing non-overlapped key-proteins; the Y-axis displays the three top-ranked drugs suggested as common treatments for both diseases.

Docking analysis was additionally performed against selected disease-associated genes related to T2D (IL6, JUN, MYC, FOS, CDC42) and HT (HSP90AA1, PTPRC, CD8A, CDC42, CD74). Although several of these proteins were not part of the defined sKG set, predicted binding affinities ≤ −7.0 kcal/mol were observed for multiple protein–ligand combinations (**Figure 10B**). These findings represent computational interaction predictions and do not establish inhibitory activity or therapeutic efficacy.

### Evaluation of pharmacokinetic parameters and toxicological profile

ADME/T tests are critical to decide if a molecule could be a safe, effective drug. Drug-likeness examines its chemical/physical traits. Good oral efficacy usually means human intestinal absorption (HIA) > 30%[36,37]. In this analysis, all drug compounds-except Naringin-exhibited high HIA scores (≥70%), suggesting efficient intestinal absorption. Caco-2 permeability is used to predict gastrointestinal absorption, where values equal to or greater than 5.150 log cm/s are considered indicative of suitable permeability[36,38]. Experimental results show strong gastrointestinal absorption for all top-ranked compounds. BBB permeability was assessed using logBB, calculated as log₁₀ (C_brain/C_blood). A logBB > 0.3 indicates effective brain penetration, while values < 0 suggest poor BBB permeability[39]. None of the proposed drug candidates showed effective BBB penetration, since all logBB values were < 0.3 (**Table 4**). The volume of distribution at steady state (VDss) quantifies the extent to which a drug disperses into body tissues. A volume of distribution exceeding 2.810 L/kg is regarded as high[37], and our molecules demonstrated favorable VDss values, suggesting extensive tissue distribution. The data suggest partial CNS penetration. Since cytochrome P450 is crucial for drug breakdown, its interactions with these compounds are especially relevant [40]. They showed good clearance (1–5 L/h), low accumulation risk, no skin sensitization, and, except for Wedelolactone, passed the AMES test without mutagenicity. Overall, their pharmacokinetics and toxicity profiles are favorable, strongly supporting their repurposing potential for T2D and HT.

**Table 4.** ADME/T profile of the top 05 drugs.

| Compounds | Absorption |  |  | Distribution |  | Metabolism (CYP450) |  |  |  | Excretion | Toxicity |  |  |
| --- | --- | --- | --- | --- | --- | --- | --- | --- | --- | --- | --- | --- | --- |
|  | Caco2 | HIA (%) | LogS | BBB | VDss | CYP1A2 | CYP2C19 | CYP2C9 | CYP2D6 | TC | AMES | Skin sensitization | Carcinogenicity |
|  |  |  |  | (Permeability) |  |  |  |  |  |  |  |  |  |
| Gliquidone | 0.636 | 69.855 | -4.539 | -0.942 | -0.152 | no | no | yes | no | 0.379 | no | no | no |
| Oleanolic acid | 1.168 | 99.558 | -3.261 | -0.143 | -1.009 | no | no | no | no | -0.081 | no | no | no |
| Glipizide | 0.582 | 63.098 | -3.414 | -1.043 | -0.249 | no | no | yes | no | 1.125 | no | no | no |
| Naringin | -0.658 | 25.796 | -2.919 | -1.6 | 0.619 | no | no | no | no | 0.318 | no | no | no |
| Wedelolactone | -0.23 | 93.753 | -3.277 | -1.35 | 0.129 | yes | no | yes | no | 0.641 | yes | no | yes |

### Drug-likeness properties

Following molecular docking simulations, the top-performing compounds-Gliquidone, Oleanolic acid, Glipizide, Naringin, and Wedelolactone subsequently assessed for their drug-likeness properties (**Table 5**). All compounds exhibited robust pharmacokinetic profiles, adhering closely to Lipinski’s Rule of Five, except Naringin, which showed some deviations. All these drugs demonstrate strong potential for repurposing.

**Table 5.** Drug-likeness profile of top 5 drugs.

| Compounds | Molecular weight | Log P | Water solubility (log mol/L) | H-bond Acceptor (HBA) | H-bond donor (HBD) | Polar Surface area (Å<sup>2</sup>) | No of rotatable bonds | Follows | Violation |
| --- | --- | --- | --- | --- | --- | --- | --- | --- | --- |
| Gliquidone | 527.64 | 3.518 | -4.539 | 6 | 2 | 217.169 | 7 | 3 | 1 |
| Oleanolic acid | 456.71 | 7.233 | -3.261 | 2 | 2 | 201.354 | 1 | 3 | 1 |
| Glipizide | 445.54 | 2.078 | -3.414 | 6 | 3 | 181.670 | 7 | 4 | 0 |
| Naringin | 580.53 | -1.165 | -2.919 | 14 | 8 | 233.028 | 6 | 1 | 3 |
| Wedelolactone | 314.24 | 2.817 | -3.277 | 7 | 3 | 127.098 | 1 | 4 | 0 |

### Molecular dynamics (MD) analysis

Molecular dynamics (MD) simulations were carried out for 100 ns to evaluate the stability, conformational flexibility, structural compactness, and binding behavior of the Gliquidone-CDC42, Oleanolic acid-CD74, and Glipizide-RAC2 complexes (**Figure 11**). The RMSD profiles showed that all three complexes maintained well-stabilized conformations throughout the simulation, with average RMSD values of 2.68 Å, 1.15 Å, and 2.33 Å (**Figure 11A**), respectively, confirming their structural robustness. Residue-level fluctuations assessed through RMSF analysis revealed mean values of 1.52 Å, 1.98 Å, and 2.75 Å (**Figure 11B**), indicating generally stable residue dynamics with only minimal localized flexibility. Collectively, these findings reflect strong protein–ligand interactions and support the overall thermodynamic stability of the complexes. These findings indicate that Gliquidone, Oleanolic acid, and Glipizide maintain stable and efficient interactions with their respective target receptors, underscoring their promise as potential therapeutic agents for managing HT in patients with T2D.

**Figure 11.**
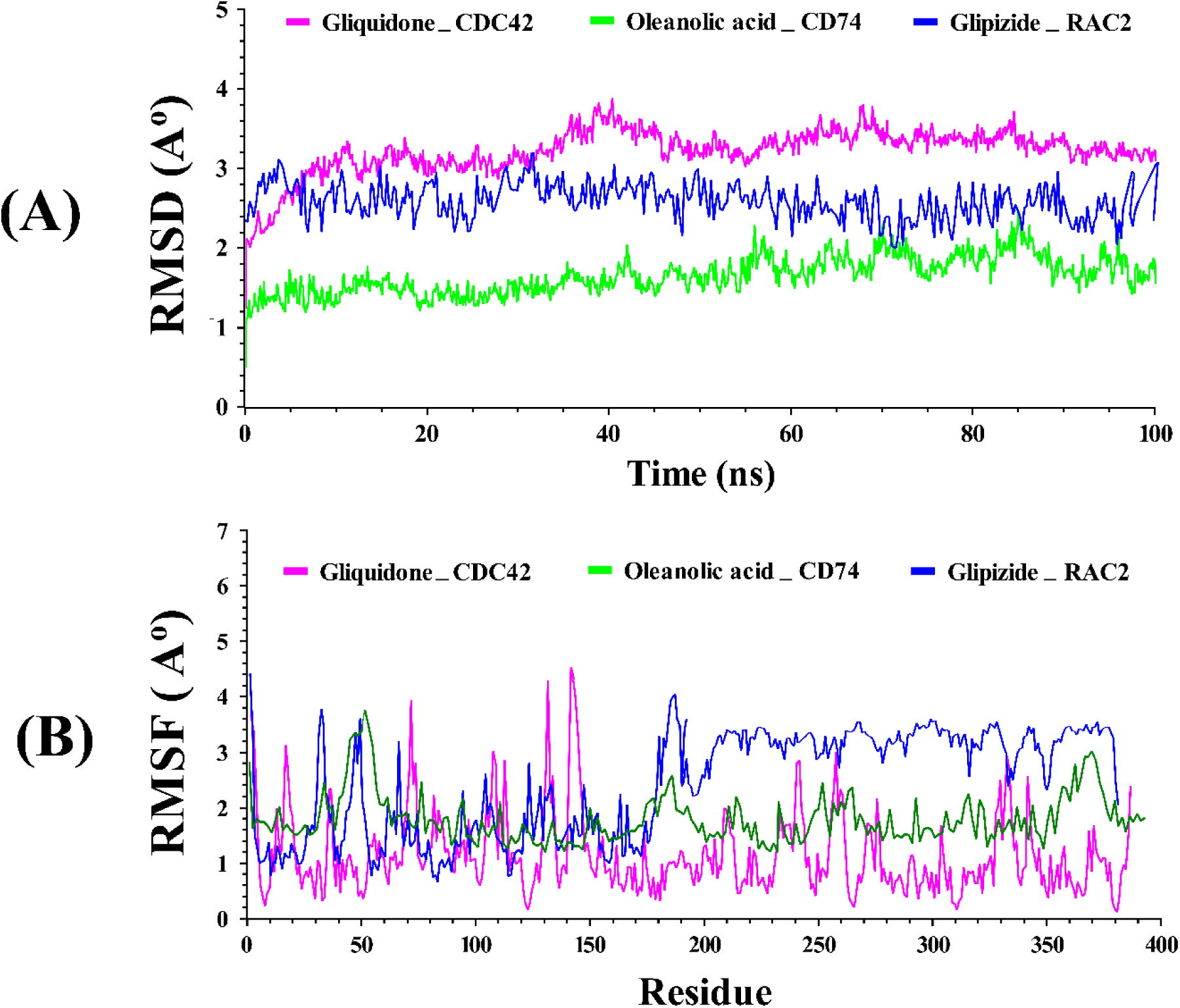
Molecular dynamics (MD) simulation outcomes for the top-ranked drug–target complexes are presented as follows: (A) root mean square deviation (RMSD), (B) root mean square fluctuation (RMSF), for the three candidate drugs over the 100-ns simulation period.

## Discussion

Hashimoto thyroiditis is an autoimmune condition that has become an important health issue but particularly in patients with underlying comorbidities, as these co-occurring conditions can significantly modify disease patterns and overall health[41,42]. Among these comorbidities, type 2 diabetes has been identified as a possible risk factor for more severe disease progression in autoimmune conditions like Hashimoto’s thyroiditis[41]. Transcriptomic profiling has emerged as a valuable approach for investigating a wide range of diseases, spanning autoimmune conditions, viral infections, cancers, and neurodegenerative disorders[43–46]. Using transcriptomic data, we identified five shared key genes -CDC42, CD74, FOS, RAC2, and YWHAB that are greatly over-represented in important Gene Ontology categories and KEGG signaling pathways, underscoring their likely role in the overlap between these two diseases. Among the sKGs we found, CDC42 is critically involved in modulating actin cytoskeleton organization, which is vital for immune cell motility and the maintenance of thyroid tissue integrity, and also in positively regulating endocytosis, which controls antigen presentation relevant to Hashimoto’s thyroiditis [47,48]. The binding of GTP and GTPase activity are key concerns of its intracellular signaling that associates with insulin resistance of T2D due to inflammatory response[47,49].Focal Adhesions and Filopodia are the main localization sites of CDC42 since it mediates interactions between cells and immune cell trafficking[50]. Additionally, it represents the converging point of the MAPK signaling pathway, which mediates pro-inflammatory or metabolic signals as part of the mechanisms that characterize the comorbidity between HT and T2D[51]. In HT, CD74 is over-expressed in thyroid cells, helping them act more like antigen-presenting cells via MHC II, which may strengthen their interaction with immune cells and fuel chronic inflammation[52,53]. In T2D, CD74 functions as a cytokine receptor for macrophage migration inhibitory factor, amplifying proinflammatory signaling and driving metabolic inflammation[54]. This dual role, further supported by its ability to enhance positive regulation of interleukin-6 production, highlights CD74 as both a mediator of thyroid autoimmunity and a contributor to systemic metabolic dysfunction, thereby linking the pathophysiology of HT and T2D[55]. RAC2, a hematopoietic-specific GTPase, regulates immune cell function through Positive regulation of neutrophil chemotaxis and Regulation of respiratory burst. These processes are essential for autoantigen-driven inflammation in Hashimoto’s thyroiditis[56–58]. RAC2 may contribute to oxidative stress and impaired insulin signaling via a MAPK-dependent pathway, thereby potentially amplifying metabolic inflammation and β-cell dysfunction[59,60]. The GTPase activity of RAC2 contributes to the remodeling of the actin cytoskeleton, both in the abnormal activation of immunity in thyroid autoimmunity and glucose homeostasis in T2D[61]. This dual role underscores RAC2 as a driver of shared inflammatory and metabolic pathways that underlie the comorbidity of HT and T2D. The FOS gene contributes to comorbidity mechanisms linking HT and T2D through immune-inflammatory and metabolic dysregulation. Its role in the MAPK signaling pathway drives pro-inflammatory cytokine expression, exacerbating pancreatic β-cell stress and thyroid autoimmunity[62].

Moreover, the IL-17 signaling pathway further amplifies chronic inflammation, promoting both thyroid tissue destruction in HT and insulin resistance in T2D[63]. At the molecular level, FOS regulates core promoter sequence-specific DNA binding, enhancing transcription of inflammatory mediators that underpin shared autoimmune and metabolic pathology[64]. The YWHAB gene, a 14-3-3 family protein, plays a regulatory role in metabolic-immune crosstalk that underlies the comorbidity between HT and T2D[65]. Through the Hippo signaling pathway, YWHAB influences cell survival and immune regulation, contributing to thyroid tissue damage and pancreatic β-cell dysfunction[66]. Its role in the cell cycle further links abnormal proliferation with chronic inflammation, aggravating both autoimmunity and metabolic imbalance[67]. At the molecular level, YWHAB may exert regulatory control through protein phosphatase interactions, with potential implications for pro-inflammatory signaling and epigenetic modulation. Putative interactions with histone deacetylases (HDACs) suggest a possible link to both thyroid autoimmunity and insulin resistance[68]. The dysregulation in these sKGs is a crucial determinant of disease progression, and thus potentiates the clinical burden of comorbidities. Regulatory network analysis identified two transcription factors, HNF4A and FOXC1, alongside two microRNAs, hsa-miR-221-3p and hsa-miR-29a-3p, as significant modulators of sKGs implicated in both HT and T2D. Notably, all sKGs were found to be targets of FOXC1 and HNF4A, indicating their central role in regulating gene expression across diverse physiological and pathological contexts. HNF4A is a nuclear receptor and transcription factor that plays a central role in regulating the genes implicated in glucose and lipid metabolism[69]. It controls the activity of genes involved in fatty acid breakdown, ketone synthesis, and fat transport, thereby affecting the body’s energy balance and overall metabolism[70]. FOXC1 is a transcription factor that modulates both inflammation and metabolism. By upregulating DNMT3B, it drives DNA hypermethylation at the CTH promoter, facilitating hepatocellular carcinoma progression and metastasis[71]. This suggests that FOXC1 may play a role in mediating the intersection of immune dysregulation and metabolic disturbances[72]. Moreover, miRNAs, including hsa-miR-221-3p and hsa-miR-29a-3p, which can regulate gene expression on a post-transcriptional level, are relevant to autoimmune thyroiditis as well as metabolic disorders. hsa-miR-221-3p is involved in the regulation of angiogenesis, apoptosis, and vascular homeostasis, which are key pathologies in the development of T2D, and consequently, the vascular disease of Hashimoto thyroiditis[73,74]. Similarly, hsa-miR-29a-3p is involved in regulating insulin signaling, glucose metabolism, along with Th1 immune responses, again highlighting its possible role in the comorbidity established between T2D and autoimmune thyroid diseases[75–77].

Analysis with immune cell infiltration patterns identified similar and immune-specific expression patterns in HT and T2D. Enrichment of memory B cells, activated memory CD4⁺ T cells, and follicular helper T cells, and a decrease in resting memory CD4⁺ T cells and M2 macrophages were observed in both conditions, suggesting a common immune-inflammatory axis. In HT, Memory B cells play a central role in the pathogenesis as they stimulate the production of autoantibodies targeting thyroid antigens[78,79]. Similarly, emerging evidence suggests that B-cell-mediated production of antibodies and proinflammatory cytokines contributes to β-cell dysfunction and the progression of insulin resistance in type 2 diabetes[80]. Thyroid tissue is the site of accumulating activated CD4 + memory T cells, which exacerbate local inflammation[81], and are also said to contribute to adipose tissue inflammation and insulin resistance elsewhere in T2D[82]. Follicular helper T cells augment germinal center responses that lead to production of autoantibodies in HT[83], and are associated with islet autoimmunity and regulation of glycemia[84]. We observed that a decline in immunoregulatory cells, specifically M2 macrophages and resting memory CD4⁺ T cells, is associated with impaired tolerance mechanisms in Hashimoto’s thyroiditis, as demonstrated by reduced M2 macrophage polarization in animal models and evidence of altered CD4⁺ T cell homeostasis in human disease[85,86]. And a deficiency in anti-inflammatory macrophage polarization in T2D. The close linkages between hub genes and these immune subsets indicate the central role of dysregulation of adaptive immunity and the loss of immune homeostasis in the development of HT-T2D comorbidity. These findings highlight hub genes as potential targets for immunomodulatory strategies that could mitigate the shared autoimmune-inflammatory pathways between endocrine autoimmunity and metabolic dysfunction. To identify potential therapeutic agents, this study performed drug repurposing guided by sKGs associated with both HT and T2D. We performed molecular docking of 247 compounds with sKGs-mediated proteins and identified five-Gliquidone, Oleanolic acid, Glipizide, Naringin, and Wedelolactone that exhibited strong binding affinities. After evaluating drug-likeness, pharmacokinetics, and toxicity, Gliquidone, Oleanolic acid, and Glipizide emerged as promising candidates for managing HT in T2D patients. In contrast, Naringin and Wedelolactone were excluded because they did not meet all pharmacokinetic criteria, which could compromise their safety and efficacy. Gliquidone (DB01251) is a second-generation sulfonylurea with the application in the treatment of T2D. It acts through suppression of the ATP-sensitive potassium (KATP) channels of the pancreatic beta-cell, resulting in cell depolarization, influx of calcium, and increased insulin release[87]. Oleanolic acid, a natural triterpenoid found in olive leaves, has been shown to have hypoglycemic effects. Hypolipidemic effects, and even cytoprotective effects against oxidative and chemotoxic stress in type 2 diabetes mellitus[88]. Glipizide, an FDA-approved second-generation sulfonylurea (DB01067), lowers blood glucose in patients with T2D by binding to pancreatic β-cell sulfonylurea receptors (SUR), closing ATP-sensitive K⁺ channels, and stimulating insulin secretion[89]. Gliquidone, Oleanolic acid, and Glipizide are established antidiabetic agents used to manage Type 2 Diabetes. Beyond their glucose-lowering effects, Gliquidone has demonstrated anti-inflammatory activity by reducing LPS-induced ERK, STAT3, and NF-κB signaling in microglial cells[90]. Oleanolic acid exhibits anti-inflammatory effects via modulation of MAPK, PI3K/Akt/NF-κB, and related pathways[91]. Glipizide has demonstrated immunomodulatory effects by inhibiting human mononuclear cell proliferation, suggesting potential beyond glycemic control[92]. These properties suggest potential therapeutic relevance in HT, a chronic autoimmune inflammatory thyroid disorder in T2D patients. In silico analyses such as molecular docking, ADMET profiling, and toxicity studies offer key preliminary. Also, MD simulations confirmed that Gliquidone, Oleanolic acid, and Glipizide form stable, low-fluctuation complexes with their respective targets, supported by favorable RMSD, RMSF profiles. But experimental confirmation is needed before moving to clinical trials. To tackle this, the cytotoxicity of the drug should initially be evaluated in vitro using host samples, with careful preparation of the culture medium to ensure accurate experimental conditions[93]. Following this, in vivo and ex vivo investigations utilizing appropriate animal models, such as Swiss albino mice, may be undertaken to comprehensively assess the pharmacological efficacy and safety of the most promising drug candidates under biologically relevant conditions [94]. Furthermore, quantitative PCR-based gene expression analysis may be utilized to validate the effects of the candidate drugs on the expression of critical molecular targets revealed through this investigation[95].

## Methodology

### Data sources and description

In order to explore shared key-genes (sKGs) associated with both T2D and HT, we have collected publicly available transcriptomics datasets from the National Center for Biotechnology Information (NCBI) database. For T2D, we downloaded two datasets with accession IDs (GSE29231[96] and GSE25724[97])generated from two different countries. Similarly, for HT, we downloaded two datasets (GSE138198[98] and GSE29315) from two different countries. One dataset per disease was used for DEG discovery, while the remaining dataset was used for independent expression consistency assessment. Details of these datasets are shown in **Table 1**.

Drug candidates were curated using a mechanism-guided filtering strategy. To explore sKGs-guided candidate compounds for hypothesis generation rather than therapeutic recommendation, we compiled an initial pool of 247 candidate molecules from published literature and curated databases. Among these, 4 candidate molecules were obtained from HT-related studies (**Table S1**), three molecules were identified through sKGs-disease interaction analysis using the DGIdb database (**Table S1**), and the remaining 240 molecules were collected from T2D-related publications (**Table S2**). This preliminary compound pool was intentionally broad and heterogeneous to capture a wide range of biologically relevant molecules for exploratory in silico screening. Subsequently, FDA-approved drugs, clinical trial candidates, or purified compounds with documented biological activity against shared key genes or enriched pathways were prioritized for downstream mechanistic evaluation.

### Identification of Differentially Expressed Genes (DEGs)

LIMMA (Linear Models for Microarray Data) is a widely used method for identifying differentially expressed genes (DEGs) between cases and control groups [99]. In this study, DEGs were used to characterize disease-associated gene expression changes. Transcriptomic datasets were normalized within each dataset using the Robust Multi-array Average (RMA) method to reduce technical variability prior to DEG identification using the LIMMA R package. Normalization and differential expression analyses were conducted separately for each dataset, and no explicit cross-study batch correction was performed because datasets for T2D and HT were analyzed separately without integration. Upregulated DEGs were selected by using the threshold at adjusted *p*-value < 0.05 and log_2_FC > 1.5, while those with an adjusted *p*-value < 0.05 and a log_2_FC < −1.5 were marked as downregulated DEGs. These thresholds have also been widely applied in different transcriptomic studies to validate the statistical and biological significance of DEGs[100–105]. Further details on the LIMMA methodology are provided in **Method S1**.

### Identification of shared DEGs (sDEGs)

We first determined upregulated and downregulated DEGs between the HT and control groups using the LIMMA approach. Similarly, we identified up- and down-regulated DEGs between the T2D and control groups using the same approach. Then we identified shared upregulated DEGs (suDEGs) between HT and T2D as well as shared downregulated DEGs (sdDEGs) between HT and T2D. Then we combined suDEGs and sdDEGs to obtain shared DEGs (sDEGs) between HT and T2D. These sDEGs represent genes that were differentially expressed in both HT and T2D relative to control samples. This overlap-based approach may preferentially identify broadly responsive genes, including immune-related genes.

### Identification of shared key genes (sKGs) from sDEGs

sKGs are identified through protein-protein interaction (PPI) networks, which map the interactions between proteins within cells to carry out their functions [106]. These sKGs were prioritized based on network topology and do not necessarily represent causal drivers of disease. To identify sKGs, a PPI network of the sDEGs was subsequently constructed using the STRING database[107]. Using the CytoHubba plugin in Cytoscape software[108], we employed 5 topological measures: Betweenness[109], Degree[110], Closeness[111], EPC, and MCC to choose the sKGs from the PPI network.

### *In-silico* validation of sKGs using independent transcriptomic datasets

To validate the differential expression patterns of the sKGs between disease and control groups, we analyzed two independent gene expression datasets from the NCBI GEO database: GSE29315 for HT and GSE25724 for T2D. The expression profiles of the core genes were visualized and validated using the ‘ggpubr’ R package[112].

### Elucidating shared pathogenetic mechanisms of sKGs

The sKGs were further analyzed to study regulatory factors, molecular functions (MFs), biological processes (BPs), cellular components (CCs), and pathways to demonstrate commonalities of processes involved in HT and T2D, as illustrated in the following sections.

### Regulatory network analysis of sKGs

A gene regulatory network captures how molecular regulators, particularly transcription factors (TFs) and microRNAs (miRNAs), coordinate to control gene expression at transcriptional and post-transcriptional levels. To identify key TFs governing the sKGs, we constructed a TF-sKGs interaction network using the JASPAR database[113] within the Network Analyst platform[114]. Similarly, miRNA-sKGs interactions were explored through the TarBase database[115] to pinpoint the most influential miRNAs regulating sKGs. These analyses provide predicted regulatory associations that may be relevant to the regulation of sKGs expression.

### Functional enrichment analysis of the sKGs

To explore the biological functions and pathways linked to the sKGs, Functional enrichment of the genes was performed through Gene Ontology (GO) term and Kyoto Encyclopedia of Genes and Genomes (KEGG) pathway analyses. We utilized web-based tools-Enrichr[35], applying Fisher’s exact test, with statistical significance assessed after Benjamini-Hochberg false discovery rate (FDR) correction.

### Link of sKGs with the immune cell infiltration analysis

The association between sKGs expression and immune cell infiltration was analyzed using CIBERSORT[116], as implemented in the IOBR R package, to estimate the relative abundance of 22 immune cell types from bulk transcriptomic data[117]. Differences between control and disease groups were compared, and Pearson correlation analysis was performed to evaluate associations between sKGs expression and immune-cell proportions after confirming normality assumptions. Only samples meeting CIBERSORT deconvolution quality criteria were included in the analysis. Statistical significance was set at p-value < 0.05. All results were visualized using ggplot2[118].

### Druggability Assessment of sKGs

Druggability analysis of the sKGs protein structures was performed using DoGSiteScorer[119] via the ProteinPlus Server [120] to identify and characterize potential ligand-binding pockets. This grid-based approach utilizes Difference-of-Gaussian filtering to identify binding cavities and evaluate structural characteristics, including volume, surface area, depth, polarity, and hydrophobicity. The calculated druggability scores were employed to rank and select potential binding pockets for further molecular docking studies. These computational predictions offer valuable structural information regarding possible ligand-binding sites.

### SKGs - guided drug repurposing

An exploratory in silico drug screening approach was used to prioritize candidate compounds that may interact with the identified sKGs proteins. Molecular docking, predicted drug-likeness, pharmacokinetic (PK) properties, toxicity estimates, and molecular dynamics (MD) simulations were used to evaluate theoretical binding behavior and physicochemical feasibility. These analyses were intended for hypothesis generation rather than therapeutic recommendations.

### Molecular docking

Molecular docking analysis was conducted as part of an exploratory in-silico screening strategy to assess potential interactions between candidate compounds and the sKGs proteins. Docking predicts the theoretical binding affinity between small molecules and protein structures, providing preliminary insight into possible molecular interactions. Structures of potential drug candidates were obtained from PubChem[121], while Receptor structures were retrieved from the Protein Data Bank (PDB)[122], AlphaFold [123], and Swiss-model[124]. The study used AutoDock tools[125]to preprocess receptors, Avogadro[126]to optimize drug compounds, and AutoDock Vina[125]for docking to determine binding affinities. Compounds were ranked based on predicted binding affinity scores (BASs) across targets to prioritize candidates for further investigation. The identified compounds should be considered preliminary candidates requiring experimental validation before any biological or clinical conclusions can be drawn. Comprehensive details of the molecular docking procedures can be found in **Method S2**.

### Evaluation of pharmacokinetic parameters and toxicological profile analysis

In drug repurposing, strong docking alone is insufficient, as compounds with poor ADME/T properties-such as limited absorption, rapid metabolism, or high toxicity-are unlikely to succeed. Therefore, evaluating ADME/T profiles is essential for selecting candidates with favorable pharmacokinetics and safety, often serving as an efficient alternative to early in-vivo or in-vitro testing[127–130]. The ADME characteristics of the candidate compounds were predicted using the ADMETlab 2.0 server[131]. Toxicity and pharmacokinetic risks were further assessed using the ProTox [132] and pkCSM[36] web tools.

### Drug-likeness properties

Drug-likeness and oral absorption potential were assessed using Lipinski’s Rule of Five alongside pkCSM [36] and ADMETlab 2.0[131]. Which predicted key physicochemical and medicinal-chemistry properties, including LogP, molecular weight, hydrogen bonding, PAINS, Brenk alerts, and synthetic accessibility. SMILES structures of all candidate compounds were retrieved from the PubChem database [83] for these evaluations.

### Molecular dynamics (MD) analysis

Molecular dynamics simulations of the top-ranked protein–ligand complexes were carried out in YASARA[133] using the AMBER14 force field to evaluate their dynamic behavior and stability. Each system was solvated with the TIP3P (Transferable Intermolecular Potential with 3 Points) water model[134], the hydrogen-bonding network was optimized, and periodic boundary conditions were applied before performing simulated-annealing minimization. The complexes were subsequently simulated for 100 ns under physiological conditions (298 K, pH 7.4, 0.9% NaCl) using a 2.5 fs [135]multiple time-step scheme regulated by the Berendsen thermostat[136,137]. Trajectories were recorded every 100 ps and analyzed using YASARA’s built-in macros[138] and SciDAVis (http://scidavis.sourceforge.net/). Binding free energies were then calculated for each snapshot using the MM-PBSA method implemented in YASARA[139], which estimates ΔG from the potential and solvation energy contributions of the receptor, ligand, and complex. In accordance with thermodynamic principles, more negative MM-PBSA values indicate stronger predicted binding affinities[140,141].

### Limitations of this study

Independent datasets were assessed in this study for HT and T2D to identify sKGs associated with both diseases, as no transcriptomic datasets were available from patients simultaneously affected by both conditions. In this study, sKGs were identified from sDEGs using PPI network analysis based on various topological parameters. However, relying solely on topological metrics in PPI networks can occasionally lead to false-positive or false-negative identification of sKGs[142]. Molecular docking-based drug screening against sKGs proteins may yield both false-positive and false-negative results. Therefore, the findings of this study require further experimental validation.

## Conclusion

The findings of this study indicate that HT and T2D share overlapping gene expression patterns and immune-related molecular features identified through comparative transcriptomic analysis. Five shared key genes (sKGs): CDC42, FOS, CD74, RAC2, and YWHAB were prioritized based on network topology and were associated with common functional annotations, regulatory interactions, and immune infiltration patterns in both conditions. Enrichment and regulatory network analyses highlighted shared biological pathways and immune-associated processes, providing descriptive insight into potential molecular commonalities. Through exploratory computational screening, gliquidone, oleanolic acid, and glipizide were prioritized as candidate compounds based on predicted interactions with the identified targets. However, these results represent in-silico predictions and do not establish therapeutic efficacy or clinical applicability. Further mechanistic validation through wet-laboratory experiments and well-designed clinical studies is required to confirm the biological and therapeutic relevance of these findings.

## Data availability

The datasets analysed in this study are freely available at the following links https://www.ncbi.nlm.nih.gov/geo/query/acc.cgi?acc=gse138198

https://www.ncbi.nlm.nih.gov/geo/query/acc.cgi?acc=GSE29315

https://www.ncbi.nlm.nih.gov/geo/query/acc.cgi?acc=GSE29231

https://www.ncbi.nlm.nih.gov/geo/query/acc.cgi?acc=GSE25724

## Author’s contribution

O.S.: Conceptualization, Data curation, Formal analysis, Investigation, Methodology, Software, Visualization, Writing - original draft. M.F.A.: Formal analysis, Methodology, Visualization. D.S.: Methodology, Visualization, A.S.: Investigation, Visualization, M.F.A.: Data curation, Formal analysis. M.S.H.: Data curation, Visualization. M.A.N.: Investigation, Methodology, M.A.L.: Writing - review & editing. M.A.: Methodology, Software, D.M.A.: Methodology, Software, M.N.H.M – Conceptualization, Methodology, Software, Supervision, Writing - review & editing.

## Acknowledgments

The authors would like to express their sincere gratitude to the researchers and institutions that made their datasets publicly available, which significantly supported this study. The authors also acknowledge the developers of the computational tools and bioinformatics resources used in this research for enabling comprehensive in silico analyses. Additionally, the authors appreciate the academic and technical support provided by Bioinformatics Lab (Dry), Department of Statistics, University of Rajshahi, Rajshahi-6205, Bangladesh.

## Supporting information

Method S1. LIMMA approach

Method S1. Molecular docking

### Supplementary Tables

Table S1. Collection of drug agents, Hashimoto’s thyroiditis, and sKGs guided drug agents

Table S2. Collection of type-2 diabetes (T2D) related candidate drug agents from published articles and other sources.

Table S3. List of DEGs for each of HT and T2D Table S4. List of sDEGs between HT and T2D

Table S5. Different topological measure scores of common share key genes (sKGs) from the PPI network based on the STRING database and Cytoscape.

Table S6. The top significantly (p-value<0.05) enriched GO terms and KEGG pathways with sKGs by the Enrichr web tool

Table S7. The top significantly (p-value<0.05) enriched GO terms and KEGG pathways with sKGs by the Enrichr web tool

Table S8. The top significantly (p-value<0.05) enriched GO terms and KEGG pathways with sKGs by the Enrichr web tool

Table S09.Residues forming each predicted binding pocket are listed below

Table S10. Docking (binding affinity) scores (kcal/mol) between the proposed target genes/proteins (receptors) and the top ordered 20 candidate drugs (out of 247).

